# Whether the ADP-ribosyltransferase activity of Ta-sro1, a noncanonical PARP protein, contributes to its function in salinity-stress tolerance?

**DOI:** 10.1101/2022.08.24.505095

**Authors:** Shu-Wei Liu, Shu-Peng Liu, Wen-Long Wang, Mei Wang, Meng Wang, Guang-Min Xia

**Affiliations:** The Key Laboratory of Plant Development and Environment Adaptation Biology, Ministry of Education, School of Life Science, Shandong University, Qingdao 266237, P. R. China; State Key Laboratory of Soil and Sustainable Agriculture, Institute of Soil Science, Chinese Academy of Sciences, Nanjing 210008, P. R. China

## Abstract

ADP-ribosylation mediated by ADP-ribosyltransferases (ARTs) is an intricate modification that regulates diverse cellular processes including DNA repair, chromatin remodeling and gene transcription responding to stresses. In addition to the canonical poly(ADP-ribose) polymerases (PARPs), plant specific SRO (Similar to RCD One) family also contain the catalytic core of the PARP domain. However, whether the PARP domains in SROs execute the ART function is still under debate. In 2014, we reported a wheat SRO, Ta-sro1, had the ADP-ribosyltransferase activity and enhanced wheat seedling growth and abiotic stress resistance, however, a recent work by Vogt et al. showed that Ta-sro1 without ADP-ribosyltransferase activity. Based on the recent progress on PARPs and SROs in relation to ADP-ribosyltransferase activity, along with our former and recent evolving results, we argued that Ta-sro1 is a non-canonical ADP-ribosyltransferase with the enzymatic activity. Although we have revealed the novel mechanism of Ta-sro1 regulate redox homeostasis and enhance salinity stress tolerance through interacting with TaSIP1, it is of interest to further clarify whether and how the enzymatic activity of Ta-sro1 responsible for the salinity tolerance of wheat. Our study raises some interesting points and caveats that helpful for understanding the research progresses and debates about the enzymatic activity of SROs.

---

*Ta-sro1* is a superior gene for trade-offs between salinity stress tolerance and yield in wheat (Wang et al., 2022b), which was identified in a highly salinity-tolerant wheat introgression cultivar Shanrong No.3 (SR3) (Liu et al., 2014). Earlier transcriptomic and proteomic analysis concluded that reactive oxygen species (ROS) homeostasis was the major biochemical basis for the salt tolerance of SR3 (Wang et al., 2009; Liu et al., 2012), which has been summarized as one of the main mechanisms for the salinity tolerance of bread wheat by some reviews (Munns and Gilliham, 2015; Wang and Xia, 2018; Yang and Guo, 2018). Several genes involved in ROS homeostasis have been decoded from SR3 (Dong et al., 2013; Wang et al., 2016, 2020; Zhao et al., 2016), among which *Ta-sro1* was identified via QTL mapping in combination with Omics analysis (Liu et al., 2014). We verified that Ta-sro1 could modulate ROS content by altering the gene expression and the activity of enzymes such as nicotinamide adenine dinucleotide phosphate-oxidase (NADPH oxidase, NOX) and alternative oxidase (AOX), *etc*. Meanwhile, we found that both SR3 and *Ta-sro1* overexpression (OE) lines showed higher genomic integrity under abiotic stress. Given that Ta-sro1 harbors a poly(ADP-ribose) polymerase (PARP) domain which has been known as an important regulator in DNA repair, we then detected the PARP activity of *in vitro* Ta-sro1 protein as well as the *Ta-sro1* transgenic (OE and RNAi) lines via a commercially available colorimetric assay (PARP Universal Colorimetric Assay Kit, Trevigen). These results display in our previous publication entitled ‘A wheat *SIMILAR TO RCD-ONE* gene enhances seedling growth and abiotic stress resistance by modulating redox homeostasis and maintaining genomic integrity’ (Liu et al., 2014). Considering the important value of *Ta-sro1* in salinity tolerance breeding, we have been devoted to exploring its regulating mechanisms. A novel mechanism of Ta-sro1 regulating redox homeostasis of SR3 has been elucidated in our recent publication ‘TaSRO1 plays a dual role in suppressing TaSIP1 to fine tune mitochondrial retrograde signaling and enhance salinity stress tolerance’ (Wang et al., 2022b). It is shown that the interaction between Ta-sro1 with a transmembrane domain-containing NAC transcription factor TaSIP1 (the key TF of mitochondrial retrograde signaling in wheat) in the cytoplasm, arrests more TaSIP1 on the endoplasmic reticulum, and in the nucleus, attenuates the trans-activation activity of TaSIP1, thus reducing the TaSIP1-mediated activation of *TaAOX1s*, avoiding the detrimental effect of a prolonged or excessive episode of mitochondria retrograde signaling on wheat development under stressful conditions and enhancing the yield in saline-alkali soil (Wang et al., 2022b). However, it is of interest to further clarify whether the SR3 salinity tolerance was also caused by elevated Ta-sro1’s poly(ADP-ribose) polymerase activity, and meanwhile this elucidation is also a reply to the query and comments from Vogt et al (2022).

Based on the recent progress on PARPs and SROs in relation to ADP-ribosyltransferase activity, along with our recently evolving results, we argued that the crystal structure of only the PARP domain and the variation of canonical His-Tyr-Glu are not enough to rule out the ADP-ribosyltransferase activity of Ta-sro1, particularly considering that Ta-sro1 harbors at least four intrinsically disordered regions (IDRs). We provided further evidence to clarify that the function of Ta-sro1, as a non-canonical ADP-ribosyltransferase, required a relatively higher concentration to show the enzymatic activity, and also showed a potential effect on mediating the cytoplasmic protein ADP-ribosylation, which has been widely discovered in animals but barely reported in plants (Vyas et al., 2013).

## Are SROs additional candidate ADP-ribosyltransferases in plants?

ADP-ribosylation is an intricate and versatile modification happening not only on proteins, but also on nucleic acids and even metabolites, that regulates diverse cellular processes including DNA repair, chromatin remodeling and gene transcription responding to biotic and abiotic stresses (Lüscher et al., 2021). ADP-ribosylation is mediated by ADP-ribosyltransferases (ARTs), that catalyzes the transfer of ADP-ribose (ADPR) from nicotinamide adenine dinucleotide (NAD^+^) onto substrates and generates mono(ADP-ribosyl)ation (MAR) or poly(ADP-ribosyl)ation (PAR) modifications (Feijs et al., 2013; Cohen and Chang, 2018). The ART superfamily consists of 23 families, which can be divided into three clades, among which the well-known PARP family proteins belong to the clade 1 (ARTD clade). PARP proteins have been found in all eukaryotes except yeast, and 17 PARP genes have been identified in the human genome (Otto et al., 2005; Perina et al., 2014). By contrast, the model plant Arabidopsis possesses just three PARPs; of them, AtPARP1 and AtPARP3 are similar to human HsPARP1, whereas AtPARP2 is similar to human PARP3. In addition to the three canonical PARPs, members of a plant specific SRO (Similar to RCD One) family also contain the catalytic core of the PARP domain together with a C-terminal RST domain and some members with an N-terminal WWE domain, which can interact with diversified proteins to regulate development and stress resistance (Jaspers et al., 2010; You et al., 2013; Qin et al., 2021). The SRO family of Arabidopsis comprises 6 members, AtRCD1 (Radical-induced Cell Death1) and its homologs AtSRO1 through AtSRO5. However, whether the PARP domains in SROs execute the ART function is still under debate. On one hand, SROs were suggested to have no ADP-ribosyltransferase activity, according to the bioinformatic analysis of the PARP domain folding structure, and also biochemical assay of AtRCD1 (Jaspers et al., 2010; Wirthmueller et al., 2018). On the other hand, we first reported that a wheat SRO, Ta-sro1, showed the ADP-ribosyltransferase activity (Liu et al., 2014), which was doubted in the letter of Vogt et al (2022). Recently, Arabidopsis AtSRO2 was also found as an ART to MARylate plant immune regulators SZF1/SZF2 at multiple Asp and Glu residues (Kong et al., 2021). In addition, it has been reported that, in contrast to pharmacological inhibition of PARPs that modify plant responses to stress and also plant growth, the knockout lines of single, double, and triple *parps* all showed no obvious alteration in development and also responses to abiotic and biotic stresses; moreover, protein PARylation was not abolished, but even increased in the *parp* triple mutant, indicating the presence of additional proteins, particularly SROs, with PARP activity in Arabidopsis (Rissel et al., 2017). Although studies on PARP (PARP-containing) proteins in plants are far lagged behind those in animals, increasing researchers with different technical backgrounds are participating in the study of this area, together to verify whether SROs are involved in ADP-ribosylation.

## His-Tyr-Glu triad in PARP domain should not be enough to indicate the ADP-riboysltransferase activity of ARTs

Vogt et al. (2022) presented a structure of the PARP domain of Ta-sro1 at 2.1 Å resolution and pointed out that Ta-sro1 should not bind NAD^+^ and is unlikely to be an ART, as the classical catalytic triad His-Tyr-Glu (H-Y-E) was replaced by Leu-His-His (L-H-H) in Ta-sro1. It was proposed that the conserved H-Y-E motif in the catalytic domains is essential for the activity of human PARPs, in which the His and Tyr residues are required for the binding of NAD^+^ and the Glu is a key catalytic residue that catalyzes the transfer of ADP-ribose to the acceptor site (Papini et al., 1989; Marsischky et al., 1995). However, recent studies revealed that human HsPARP9, which harbors a ‘Q-Y-T’ triad and was thought inactive previously, showed the ADP-riboysltransferase activity when it interacted with a histone E3 ligase Dtx3L forming the Dtx3L/PARP9 heterodimer (Yang et al., 2017). Furthermore, Arabidopsis AtSRO2, which does not harbor the ‘H-Y-E’ but a ‘Y-H-N’ triad, could directly, not requiring a partner, shows the NAD^+^-dependent ADP-riboysltransferase activity (Kong et al., 2021). Vogt et al. (2022) suggested that the remaining catalytic activity of AtSRO2 is due to the swap of His with Tyr in the PARP domain. However, the properties of Tyr and His are very different, and therefore their swap may influence the structure of the PARP catalytic domain dramatically, making the explanation unacceptable. Actually, the ‘H-Y-E’ triad is not the sole indicator of the ADP-riboysltransferase activity for PARP proteins, and for example, the D-loop also affects the activity (Vyas et al., 2014). Moreover, the three clades of ARTs share a comparable overall structure consisting of a split β-sheet and two helical regions (which was harbored by Ta-sro1 as revealed by Vogt et al. (2022)), whilst the primary amino acid sequences show low conservation (Lüscher et al., 2021). More recently, it was reported that the Toll-Interleukin-1 receptor (TIR) domain without ‘H-Y-E’ triad catalyzes ADP-ribosylation of adenosine triphosphate (ATP) and ADP-ribose via ADPR polymerase-like and NADase activity (Jia et al., 2022). As SROs including Ta-sro1, showed lower overall sequence identities with canonical PARPs, the amino acid sites other than the catalytic triads may also affect the enzymatic activity. Collectively, we suggest that the presence of His-Tyr-Glu triad was not essential for the catalytic activity of the noncanonical PARPs, including Ta-sro1.

## Crystal structure of only PARP domain cannot reflect the whole property of Ta-sro1

Vogt et al. (2022) depicted the crystal structure of the PARP domain of Ta-sro1. As it is shown in Figure 1D of Vogt et al. (2022), Ta-sro1 is a non-canonical PARP protein, the same as most other SRO proteins in plants, it contained an RST domain, which can interact with a variety of proteins (Jaspers et al., 2009), and the WWE domain, which can bind the poly(ADP-ribose) (Vainonen et al., 2020). More importantly, similar to RCD1 in Arabidopsis and barley, Ta-sro1 also contains at least four intrinsically disordered regions (IDRs), among which two are in the flanking region of the PARP domain (Vogt et al. (2022) in Figure 1D). IDRs are structurally flexible, which can form different conformational functional states under different conditions, allowing variability in the exposed surfaces and/or recognition motifs (Covarrubias et al., 2017). As IDRs can affect the structure of Ta-sro1, they may influence its enzymatic activity, and therefore, the activity of Ta-sro1 may not be well explained by the structure of only the PARP domain. Moreover, it has been reported that IDRs of AtRCD1/HvRCD1 are able to present a range of conformational states upon binding to its interacting partners (Covarrubias et al., 2020). In recent years, we also identified a great variety of proteins interacting with Ta-sro1, some of which were mediated by WWE and PARP domain respectively, such as TaSIP2, through the IDRs (Figure 1), but not mediated by the well-known RST domain (Jaspers et al., 2009). Meanwhile, in the human PARP family, recent studies reveal that the interaction between Dtx3L and PARP9, or between HPF1 and PARP1 or PARP2 has a significant effect on the enzymatic activity. These clues suggest a second layer of the influence by IDRs, with respect to the protein interaction, on the ADP-riboysltransferase activity for Ta-sro1. Collectively, the crystal structure of only the PARP domain is unlikely to reflect the whole property of Ta-sro1, particularly considering the existence of IDRs.

**Figure 1.**
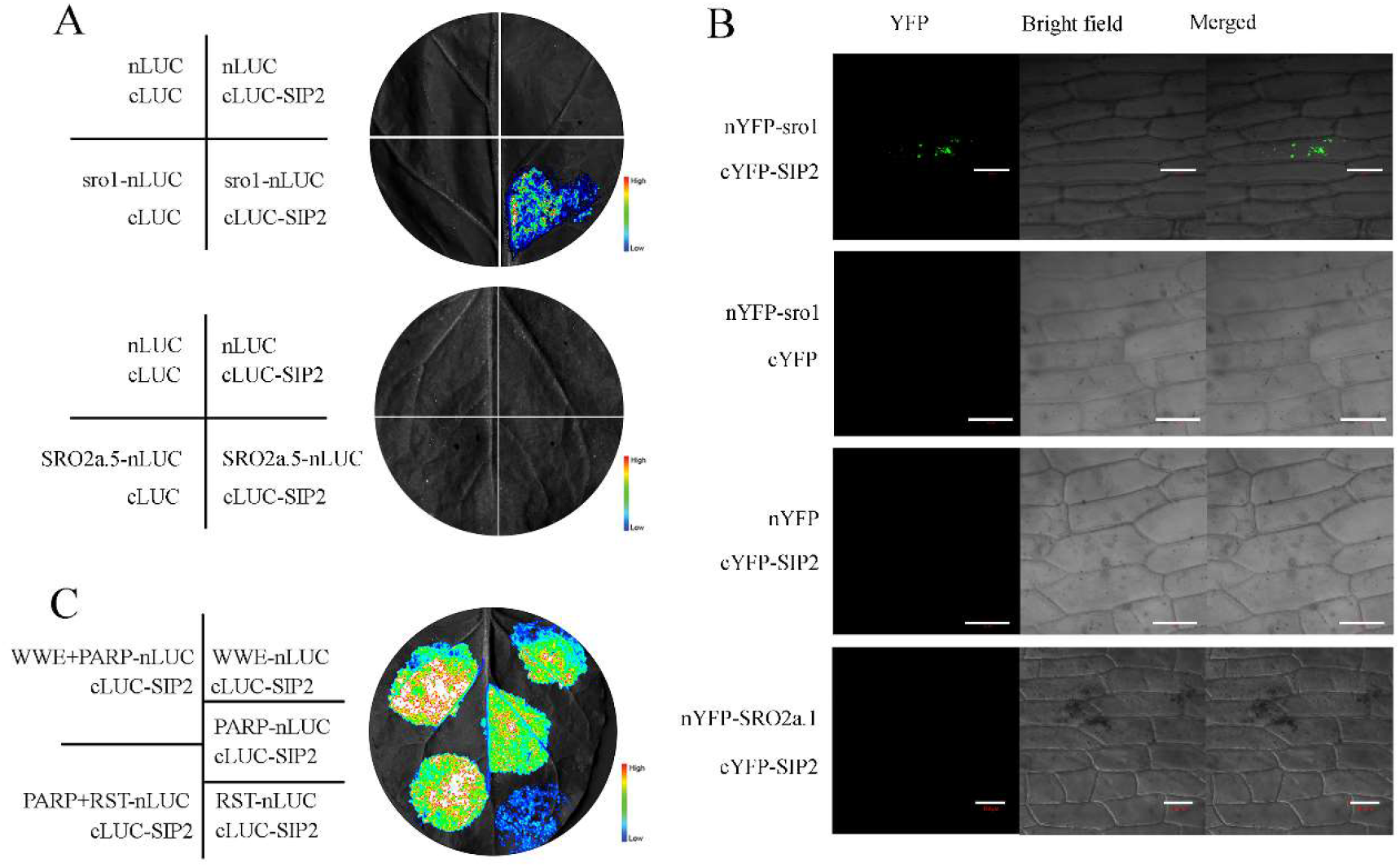
Ta-sro1 interacts with TaSIP2. (A) Split luciferase complementation (SLC) assay verified the interaction between Ta-sro1 and TaSIP2 in *Nicotiana benthamiana* leaves. TaSRO2a.1 (TraesCS1D02G199200.1), a wheat SRO protein belonging to the II-a subfamily, was used for negative control. (B) Bimolecular fluorescence complementation (BiFC) assay verified the interaction between Ta-sro1 and TaSIP2 in onion epidermal cells. TaSRO2a.1 was used for negative control. Scale bars are 100 μm. (C) SLC assay indicated that the WWE and PARP domain but not the RST domain of Ta-sro1 interacted with TaSIP2.

## Ta-sro1 itself exerts lower enzymatic activity compared to the canonical PARPs

As mentioned above, in our previous publication (Liu et al., 2014), the direct biochemical evidence for the PARP activity of Ta-sro1 was obtained from the measurements of *in vitro* Ta-sro1 protein as well as the *Ta-sro1* transgenic (OE and RNAi) lines via a commercially available colorimetric assay (PARP Universal Colorimetric Assay Kit, Trevigen). Using the same expressing plasmid we shared, Vogt et al. (2022) obtained the Ta-sro1 protein and repeated the *in vitro* measurement with the same kit, but found no obvious activity, whilst the activity of human HsPARP1 was able to be detected. We carefully looked over our original experimental records and found that 5 μg of Ta-sro1 protein per well was used in our previous experiment, while 0.5 μg of Ta-sro1 protein was used in Vogt et al. (2022). Note that there is no clearly recommended dosage for the purified protein in the manufacturer’s protocol of this kit, but a dosage of ‘at least 20 μg of protein per well’ is recommended for detecting ‘PARP activity in cell extracts’. Nevertheless, these two results attracted us to note that the enzymatic activity of Ta-sro1 is lower than that of the canonical HsPARP1. Nowadays, new approaches to detect the ADP-riboysltransferase activity are emerging and successfully applied in plants (Kong et al., 2021). To further evaluate the enzymatic activity of Ta-sro1, we performed the *in vitro* auto-ADP-ribosylation assay, which was also done in Vogt et al. (2022). Contrary to those reported by Vogt et al. (2022), we proved that Ta-sro1 showed auto-ADP-ribosylation activity. Our results indicated that when <1 μg MBP-Ta-sro1 recombinant proteins were incubated with biotin labelled NAD^+^, no auto-ADP-ribosylation was detected, however, when more than 2 μg MBP-Ta-sro1 recombination proteins were incubated with biotin labelled NAD^+^, ADP-ribosylated Ta-sro1 proteins were detected by immunoblotting using HRP-conjugated streptavidin for biotinylated NAD^+^ (Figure 2A); by contrast, ADP-ribosylated AtPARP2 proteins were detected when 0.5 μg GST-AtPARP2 recombination proteins were incubated with biotin labelled NAD^+^ (Figure 2A).

An intriguing issue is whether and how this low enzymatic activity of Ta-sro1 functions *in planta*. Our results revealed that the enhancement of seed germination in *Ta-sro1* transgenic wheat lines under salinity stress can be suppressed by PARP inhibitor 3-methoxybenzamide (3-MB) (Figure 2B), suggesting that the enzymatic activity of Ta-sro1 contributes to the function of Ta-sro1 *in planta*. With respect to how this low enzymatic activity of Ta-sro1 functions *in planta*, one possible warrant, as mentioned above, is that SROs (Jaspers et al., 2009) including Ta-sro1 (as shown in Figure 1) could interact with multiple proteins. Screening the interactor, which is able to enhance the enzymatic activity of Ta-sro1, is clearly warranted in future research. In recent years, IDRs have been shown to be a prevalent property of proteins that induced the formation of self-assembled, membrane-less organelles through liquid-liquid phase separation (LLPS) (Emenecker et al., 2020; Huang et al., 2021). AtRCD1 was found to bind and recruit Photoregulatory Protein Kinases (PPKs) to non-membranous compartments, so-called “nuclear bodies” via IDRs (Vainonen et al., 2020). Intriguingly, our results revealed that cytoplastic interaction/colocalization of Ta-sro1 with its interactors was in a punctate pattern (Figure 1B and Wang et al., 2022b), which has also been discovered in LLPS (Jiang et al., 2022). We speculate that Ta-sro1 and its’ interacting proteins could be condensate into a high concentration through LLPS, and thus maybe exert its enzymatic activity. Intriguingly, similar to those observed in AtRCD1 (Shapiguzov et al., 2019), Ta-sro1 could also aggregate as oligomers through the formation of intermolecular disulfide bonds between cysteine residues in the IDRs (Figure 3), which may help to increase the local concentration of Ta-sro1 proteins and thus enhance its enzymatic activity.

**Figure 2.**
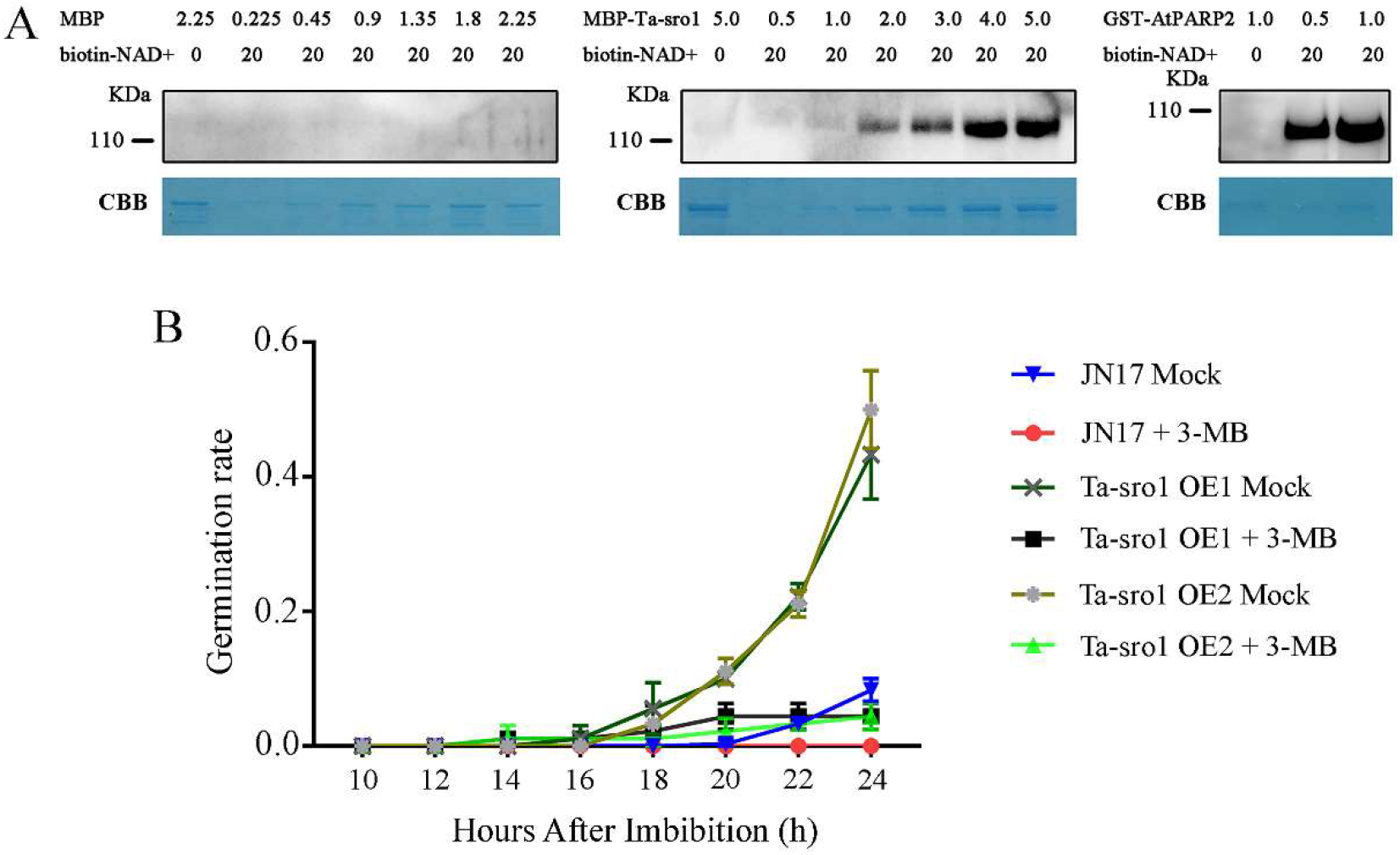
The ADP-riboysltransferase activity of Ta-sro1 and its contributions *in planta*. (A) *In vitro* ADP-ribosylation assay indicated that Ta-sro1 showed auto-ADP-ribosylation activity. To assess the auto-ADP-ribosylation activity of Ta-sro1, MBP, MBP-Ta-sro1 and GST-AtPARP2 were incubated with biotin-NAD^+^ at 25°C for 2 h and the ADPrylated proteins were detected by using Strep-HRP (Thermo Scientific, 21130). CBB, Coomassie Brilliant Blue stain. (B) PARP inhibitor 3-MB suppressed the enhancement of seed germination under salinity stress by Ta-sro1. OE1 and OE2 were two transgenic wheat lines constitutively overexpressing *Ta-sro1* in wheat cultivar JN17.

**Figure 3.**
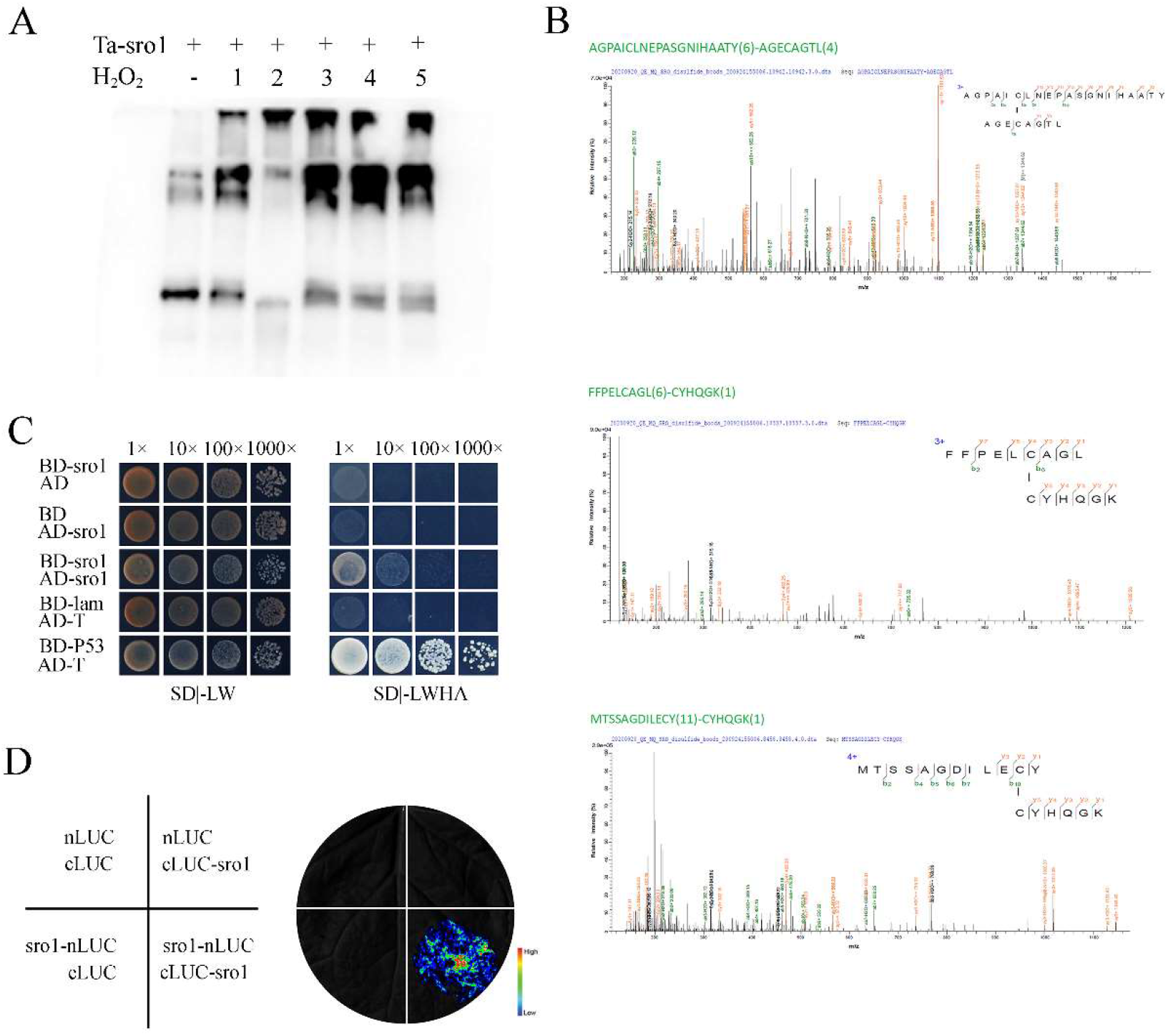
Ta-sro1 can aggregate as oligomers through the formation of intermolecular disulfide bonds. (A) SDS-PAGE analysis indicated that Ta-sro1 could aggregate as oligomers, especially under oxidative stress treatment (1-5 mM H_2_O_2_). (B) LC-MS analysis identified the oxidative Cys residues responsible for the formation of intermolecular disulfide bonds. (C) Yeast two-hybrid assay verified Ta-sro1 could form a homodimer in yeast. (D) SLC assay indicated that Ta-sro1 could form a homodimer in *N. benthamiana* leaves.

## SROs may be the potential cytoplasmic ARTs in plants

In human, several PARP proteins localize to a significant extent in the cytoplasm (Vyas et al., 2013), and the critical role of cytoplasmic ADP-ribosylation is emergingly reported (Wang et al., 2022a). In contrast, Arabidopsis possesses just two functional PARPs, AtPARP1 and AtPARP2, both of which are predominantly localized in the nucleus; whether there are cytoplasmic PARPs in plants is still unknown. Vogt et al. (2022) highlighted the role of SROs as a transcriptional co-regulator, largely due to previous discoveries that SROs interact with multiple transcription factors (TFs) in the nucleus and regulate downstream targets (Jaspers et al., 2009). However, it has been reported that AtRCD1 interacts with AtSOS1 in the cytoplasm, under the condition of high salt or oxidative stress (Katiyar-Agarwal et al., 2006). In our recent studies, Ta-sro1 was found localized both in nucleus and cytoplasm, in which it could interact with not only TFs (Figure 1B and Wang et al., 2022b) but also other type of protein (Figure 4). For example, Ta-sro1 interacts with TaSIP1 in the cytoplasm as described above (Wang et al., 2022b). In addition, the NOX (TaRboh, TraesCS3D02G279900), a membrane-bound enzyme complex, was also able to interact with Ta-sro1 (Figure 4). The interaction between Ta-sro1 and TaRboh was further proved through bimolecular fluorescence complementation (BiFC) and split luciferase complementation (SLC) assays (Figure 4), in agreement with our previous results that the NOX activity is obviously affected by *Ta-sro1* (Liu et al., 2014). These discoveries suggest that Ta-sro1 not only functions as a transcriptional co-regulator in the nucleus, but also influences the shuttling and activity of its interacting proteins in the cytoplasm, as a potential cytoplasmic ADP-ribosyltransferase.

**Figure 4.**
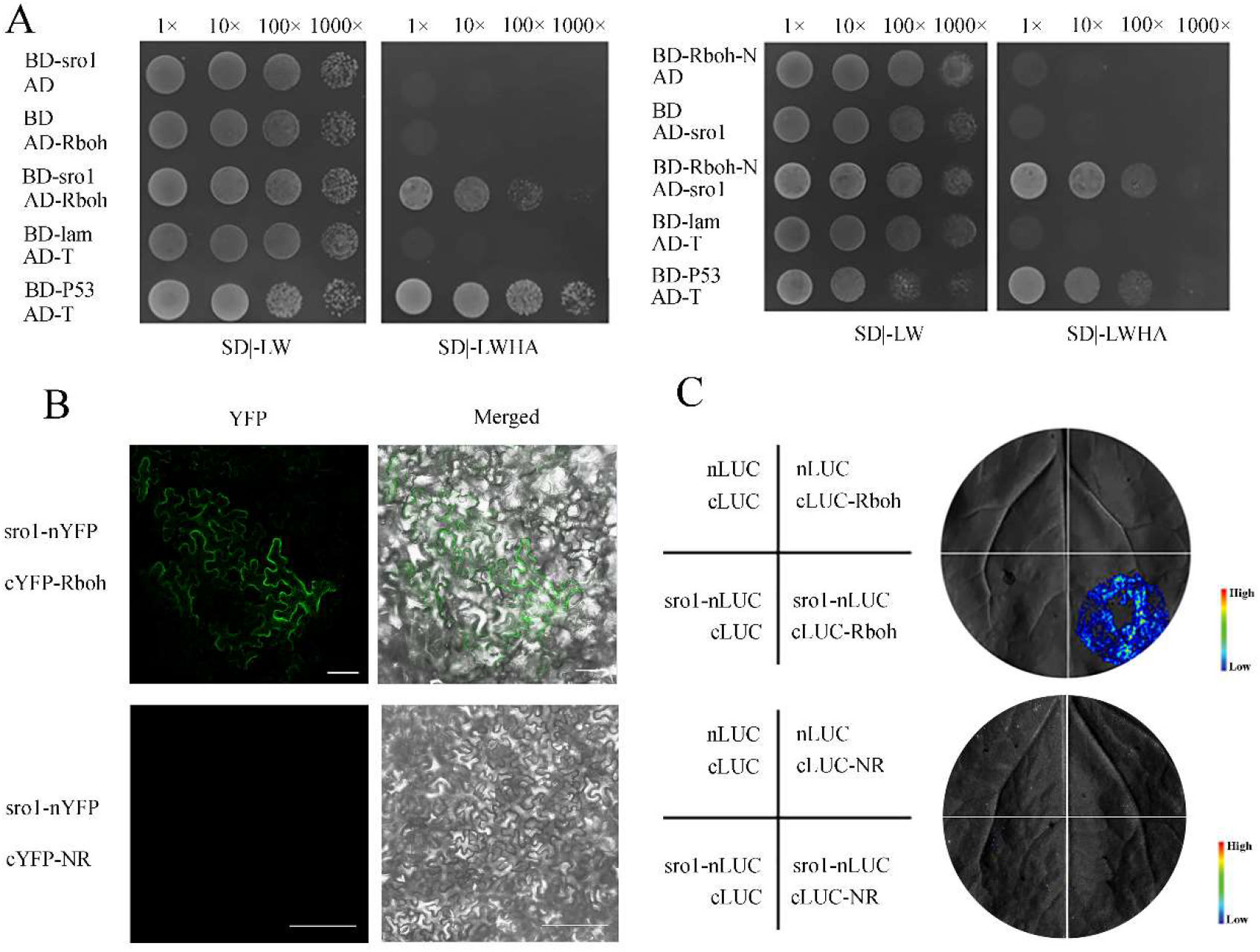
Ta-sro1 interacts with TaRboh. (A) The intracellular portion of TaRboh interacts with Ta-sro1 in yeast cells. (B) BiFC assay verified the interaction between TaRboh and Ta-sro1 in *N. benthamiana*. TaNR (TraesCS6B02G024900.1), a wheat nitrate reductase, was used for negative control. Scale bars are 100 μm. (C) SLC assay verified the interaction between TaRboh and Ta-sro1 in *N. benthamiana*. TaNR was used for negative control.

As reviewed by Suskiewicz et al. (2020), the research progress of human PARPs is in company with ‘expect the unexpected’. Similarly, since the discovery of the first SRO member AtRCD1 (Belles-Boix et al., 2000; Overmyer et al., 2000), the understanding of the characteristics and functions of SRO family members is gradually refreshed with unexpected results. Members of the SRO family were formerly considered to be nuclear localized, however, AtRCD1 and recently Ta-sro1 were found not only in the nucleus but also in the cytoplasm. The SROs were previously thought to have no ADP-ribosyltransferase activity, however, both Ta-sro1 and AtSRO2 were proved to possess ADP-ribosyltransferase activity (Liu et al., 2014; Kong et al., 2021). Recently, LLPS mediated by the IDRs was found to be involved in the function of AtRCD1 (Vainonen et al., 2020), which may enable us to further understand the functions of these noncanonical PARP proteins. With the development of mass spectrometry, gene editing and other new technologies, the enzymic activity of SROs and the underlying mechanism for regulating their interacting proteins will be further clarified at both biochemical and genetical levels, which will give us a more comprehensive understanding of this plant-specific, noncanonical PARP protein family.

## Materials and methods

### Plant materials and growth conditions

The *pUbi:Ta-sro1* transgenic wheat lines are in Jinan (JN17) background. The materials used for salt tolerance analysis were grown in an artificial climate chamber at a relative humidity of 70% and a 16 h light (26°C)/8 h dark (20°C) photoperiod, with a light intensity of 3000 lux. For salt tolerance analysis, wheat seeds of both WT and transgenic lines were sown in Petri dishes with 5 mL solution containing 200 mM NaCl and 5 mM 3-MB. The germination rate was counted every 2 hours.

### Split luciferase complementation assay (SLC)

The full-length coding sequence (CDS) or truncated derivatives were fused with the N- or C-terminal region of luciferase gene. The agrobacteria strain GV3101 harboring each construct was infiltrated into *N. benthamiana* leaves. After 3 days, the leaves were incubated into 1 mM luciferin for 5 minutes and luminescent signals were detected by Tanon-5200 Chemiluminescent Imaging System (Tanon Science and Technology, Shanghai, China).

### Yeast two hybrid assay

For yeast two hybrid assay, coding sequences of each gene cloned into vectors pGBKT7 or pGADT7. The freshly extracted plasmids were transformed into yeast strain Y2H gold by using a yeast transformation kit (TaKaRa, Beijing, China) according to the manufacturer’s protocol. After growning on SD/-Leu Trp medium, the yeast cells were screened on SD/-Leu Trp His Ade medium.

### Bimolecular fluorescence complementation assay (BiFC)

Coding sequences of each gene were amplified by PCR and cloned into the vector pSPYNE-N173 or pSPYCE (MR), respectively. Different combinations of plasmids were delivered into onion epidermal cells by particle bombardment. After 12 hours of incubation, the fluorescence microscopy was performed using a confocal laser scanning microscope (ZEISS LSM 900; ZEISS, Oberkochen, Germany).

### *In vitro* oxidation assay

His-tagged Ta-sro1 protein was purified from *E. coli* strain BL21 (DE3) and then treated with different concentrations of H_2_O_2_ in oxidation buffer (50 mM MES-KOH, pH 6.5, 100 mM NaCl, 1% Triton X-100) at room temperature for 15 min. The proteins were precipitated by adding one volume of TCA/acetone at −20°C for 20 min. The protein samples were centrifuged at 15,000 ×*g* for 20 min. The pellets were washed three times with 50% acetone and dissolved in 10 μL loading buffer without DTT and subjected to separate on 10% SDS-PAGE. All of forms of Ta-sro1 were detected with anti-His antibody (Abcam, Cambridge, UK).

### Identification of oxidative Cys by mass spectrometry

Oxidative samples were transferred into 10 KDa ultrafiltration centrifuge tubes and centrifuged at 12,000 ×*g* for 15 min. Then the 200 μL UA buffer was added and centrifuged at 12,000 ×*g* for 15 min. After centrifugation, 100 μL NH4HCO3 buffer was added and centrifuged at 14,000 ×*g* for 10 min. Trypsin and Chymotrypsin were added, shaking at 600 rpm for 1 min, and digested at 37°C for 16-18 h. The peptides were desalted with C18 StageTip. The dried peptides were re-dissolved with 0.1% FA solution for LC-MS analysis (Q-Exactive, Thermo Scientific, Waltham, MA, USA).

### ADP-ribosylation assay

For *in vitro* ADP-ribosylation assay, MBP and MBP-Ta-sro1 recombination proteins were expressed in the *E. coli* BL21 (DE3) strain and purified with amylose agarose beads (New England Biolabs, Ipswich, MA, USA) according to the manufacturer’s protocol. Different amount of Ta-sro1 proteins were incubated in 20 μL of PARylation buffer (50 mM Tris-Cl, pH 8.0, 50 mM NaCl, 10 MgCl2, 20 μM biotin-NAD^+^, 1×active DNA) at 25°C for 2 h. ADP-ribosylated proteins were separated by 10% SDS-PAGE and detected by using Strep-HRP (Thermo Scientific, Waltham, MA, USA).

## Author’s contribution

Shu-Wei Liu and Guang-Min Xia conceived the study, designed experiments, and wrote the manuscript. Shu-Peng Liu and Wen-Long Wang performed yeast two hybrid assay and *in vitro* ADP-ribosylation assay. Shu-Peng Liu performed the experiments for protein interaction verification, Ta-sro1 oligomerization, subcellular localization and PARP inhibitor treatment. Mei Wang provided the original data for PARP activity assay. Meng Wang reviewed and revised the manuscript and provided critical feedbacks.

